# Comparable affinity of RabGDIα to GTP- and GDP-bound forms of Rab7 supports a four-state transition model for Rab7 subcellular localization

**DOI:** 10.1101/287516

**Authors:** Akane Kanemitsu-Fujita, So Morishita, Svend Kjaer, Mitsunori Fukuda, Giampietro Schiavo, Takeshi Nakamura

**Author notes:** Correspondence and requests for materials should be addressed to T.N.

## Abstract

Endolysosomal system is linked to almost all aspects of a cell’s life and diseases, and Rab7 occupies a critical node in this endocytic degradation pathway. However, there have been conflicting theories about the exact role of Rab7 in membrane trafficking, since some researchers report that Rab7 regulates the trafficking from early to late endosomes, while others have reported its functions in trafficking between late endosomes to lysosomes. In the present study, we have revisited this issue from a new perspective. In COS-7 cells, the GDP-bound Rab7 mutant, T22N, was found to be located on the vesicular membranes as well as in the cytoplasm. On the other hand, the GTPase-deficient Q67L mutant of Rab7 resided in cytoplasm as well as on membranes. Additionally, we found that RabGDI interacted with both GTP and GDP bound forms of Rab7 *in vitro.* Based on the results we have proposed a four-state transition model for Rab7. This four-state model correlates with our recent findings that Rab7 was initially recruited to macropinosomes in a GDP-bound inactive form and subsequently activated during endocytic maturation in EGF-stimulated COS-7 cells.

## Introduction

Endocytic processes are linked to almost all aspects of a cell’s life and diseases. The endocytic transport of proteins and lipids is initiated in the plasma membrane. Endocytic vesicles then fuse with early endosomes (EEs), which mature into late endosomes (LEs) prior to their fusion with lysosomes. By sorting, processing, recycling, storing, activating, silencing, and degrading incoming substances and receptors, endosomes are responsible for the regulation and fine-tuning of numerous pathways in the cell (1,2). Rab5 and Rab7 are of key importance for the endolysosomal system (1,3). Rab5 functions in EEs, whereas Rab7 is required for LEs and lysosomes. Although the role of Rab5 and Rab7 as crucial factors in endolysosomal transport is widely accepted, the coordination of their transition and their multiple interactions on endosomes and lysosomes is far from being completely understood (1,2,4).

Rab7 occupies a critical node in the endolysosomal pathway. It is thought that Rab7 governs the transition from EEs to LEs, their trafficking, and fusion to lysosomes. However, there have been conflicting theories about the exact role of Rab7 in membrane trafficking, as some researchers have reported that Rab7 regulates the trafficking from early to late endosomes (5,6), while others revealed that it functions in trafficking between LEs (or autophagosomes) and lysosomes (7–9) progression. In Rab7-depeleted HeLa cells, the exiting of cargos, such as EGF-EGFR complex, from the LE/MVB was blocked, resulting in the accumulation of enlarged LEs tightly packed with intraluminal bodies (ILVs) (9). This result shows that Rab7 is dispensable for the delivery of cargos to the LEs but is required for efficient fusion of the LEs to the lysosomes, at least in EGF/EGFR complex trafficking. However, the restricted role of Rab7 during LE-lysosome fusion contradicts the widely broadly accepted view that Rab7 localizes to both the LEs and lysosomes (10), because it is thought that Rab7-GDP (inactive form) is bound to GDI and thereby resides in the cytoplasm, whereas most researchers assume that the active GTP-form of Rab7 associates with membranes.

In this study, we have revisited this issue. In COS-7 cells, the GDP-bound Rab7 mutant (Rab7 T22N), was found to be located on the vesicular membranes as well as in the cytoplasm. On the other hand, the constitutively —active Rab7 Q67L mutant resided in the cytoplasm as well as on membranes. Furthermore, RabGDI interacted with both GTP and GDP bound forms of Rab7 *in vitro.* Based on these results, we have proposed a four-state transition model for Rab7 (Fig. 3B). We would like to emphasize that this four-state model correlates with one of the major findings in our recent work 11), i.e., Rab7 was initially recruited to macropinosomes in a GDP-bound inactive form and subsequently activated during maturation in EGF-stimulated COS-7 cells.

## Results

### GDP-bound Rab7 mutant T22N, resides on the vesicular membranes as well as in the cytoplasm

We first examined the subcellular localization of Rab7 T22N in COS-7 cells. According to Fig. 1B in Yasuda et al. (11), most of the exogenously-expressed Rab7 was GDP-bound. However, Fig. 1A in this study showed that Rab7 T22N was localized on vesicular structures, probably LEs and lysosomes, as well as in the cytoplasm. We could hardly detect Rab7 T22N at the cell contour, indicating that very little or no Rab7 T22N was localized to the plasma membrane (Fig. 1A).

**Fig. 1.**
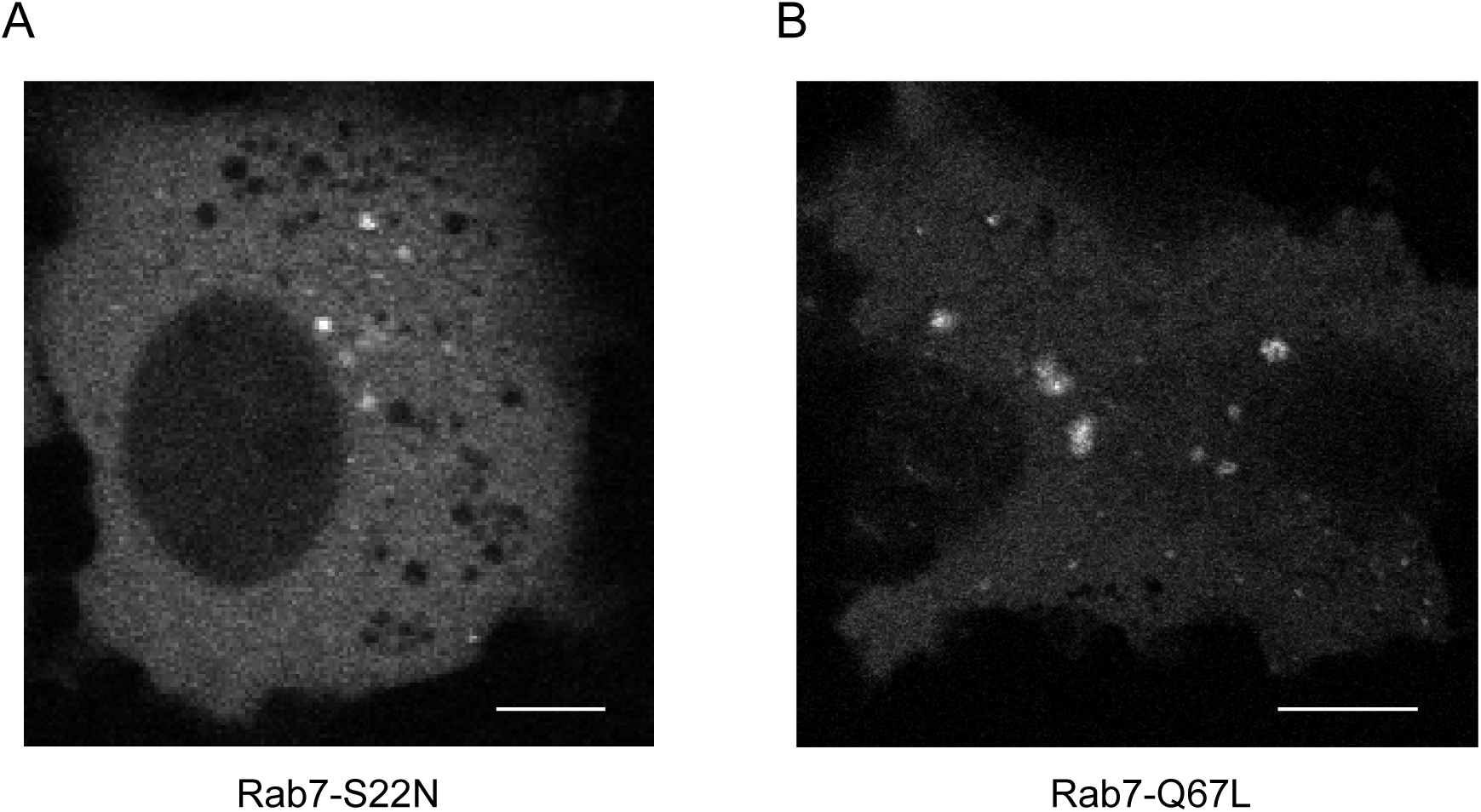
Subcellular localization of Rab7 mutants. A representative image of the subcellular distribution of (A) EGFP-Rab7-S22N and (B) EGFP-Rab7-Q67L in **COS-7** cells. Line bar, 5 µm.

### The GTP-bound Q67L mutant of Rab7 resides in the cytoplasm as well as on the membranes

Next, we examined the subcellular distribution of Rab7 Q67L in COS-7 cells (Fig. 1B). According to Fig. 1B in Yasuda et al. (11), the GTP loading of EGFP-Rab7 Q67L used here was 57%. If the two-state transition model (Fig. 3A) is correct, around 57% of EGFP-Rab7 Q67L should be in the GTP-bound form and localize to the membrane structure, and the remaining (ca. 43%) fraction, GDP-bound and residing in the cytoplasm. Our estimation of EGFP-Rab7 Q67L-expressing cells which include that shown in Fig. 1B is contradictory to this prediction. Thus, we tried to test the association of RabGDI with GTP and GDP bound forms of Rab7.

### RabGDI interacted with both GTP- and GDP-bound forms of Rab7 *in vitro.*

To clarify the nucleotide dependency of the interaction between Rab7 and RabGDI, we performed an *in vitro* binding assay with immobilized GST-RabGDIα and recombinant prenylated Rab7 loaded with GDP or GTPγS. The GTP-bound form of Rab7 binds RabGDIα with a similar binding activity as Rab7-GDP (Fig. 2). Together with the result shown in Fig. 1B, this binding assay strongly suggests that Rab7-GTP can bind RabGDIα and reside in the cytoplasm.

**Fig. 2.**
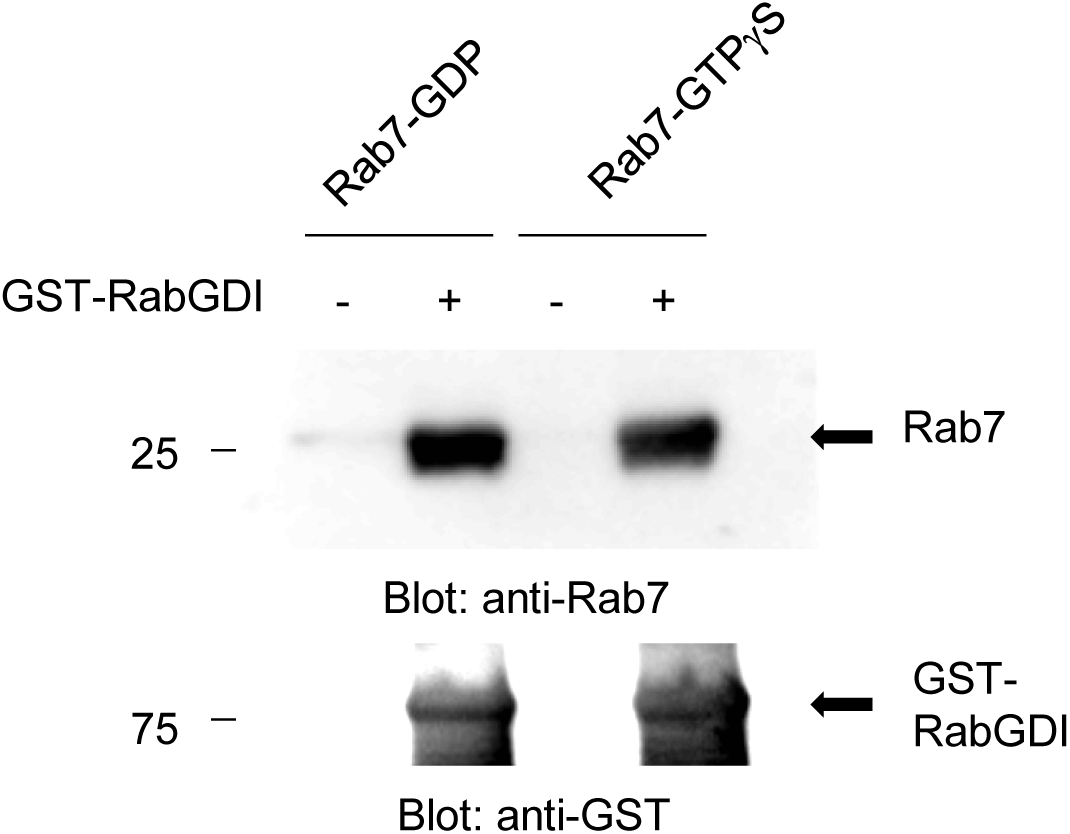
Affinity of GDP-Rab7 and GTP-Rab7 to RabGDI. The nucleotide dependency was examined by an *in vitro* GST pull-down assay. His-tagged Rab7 prenylated in insect cells and GST-RabGDI was incubated, washed, and then eluted. The elutes were immunoblotted with anti-Rab7 antibodies or anti-GST antibody, respectively.

## Discussion

In this study, we propose a four-state transition model for the subcellular localization of Rab7 (Fig. 3) based on our findings that Rab7-GDP resides on the endosomal membranes and Rab7-GTP resides in the cytoplasm. The existence of former is supported by three evidences. First, Rab7 activity of individual endosomes showed a significant variation different cells at steady-state (Fig. 2 in Yasuda et al. (11)). Second, a fraction of Rab7-T22N, which was practically regarded as Rab7-GDP, because its GTP loading was almost zero (Fig. 1B in Yasuda et al. (11)), was localized to endosomes. Third, during macropinocytosis, Rab7 activity gradually increased after initial recruitment to macropinosomes (Fig. 3 in Yasuda et al. (11)), *In vitro* binding assay showed that the GTP-bound form of prenylated Rab7 interacted with RabGDI with an affinity similar to that of as Rab7-GDP, indicating that Rab7-GTP can exist in the cytosol. For example, the previous experiment with Rab1-D44N, a Rab1 mutant that is unable to bind to RabGDI, showed that the distribution of Rab1 in the cytoplasm is completely dependent on the interaction with RabGDI (12). The possible existence of Rab7-GTP in the cytoplasm is supported by the distribution of the FRET/CFP ratio of cytoplasmic Raichu-Rab7 Q67L in several individual COS-7 cells (Kanemitsu-Fujita and Nakamura, unpublished). Compared to Raichu-Rab7 T22N, the distribution of the FRET/CFP ratio of Raichu-Rab7 Q67L, whose GTP loading was 54% (11), shifted significantly higher in the cytoplasm. This result is explainable by postulating cytoplasmic Rab7-GTP. Similarly, a weak interaction between Rab5-GDP and RabGDI has been previously reported (13). The comparison between the prevailing and largely accepted two-state model and the four-state model proposed here raises the question on how Rab7 is recruited and activated or inactivated in living cells. Our previous study (11) has partially addressed this question. Indeed, in EGF-induced macropinocytosis in COS-7 cells, Rab7 was initially recruited to macropinosomes in a GDP-bound form and subsequently activated by Mon1-Ccz1 during endosome maturation leading to a fusion with lysosomes whose Rab7 activity is independent of Mon1-Ccz1.

**Fig. 3.**
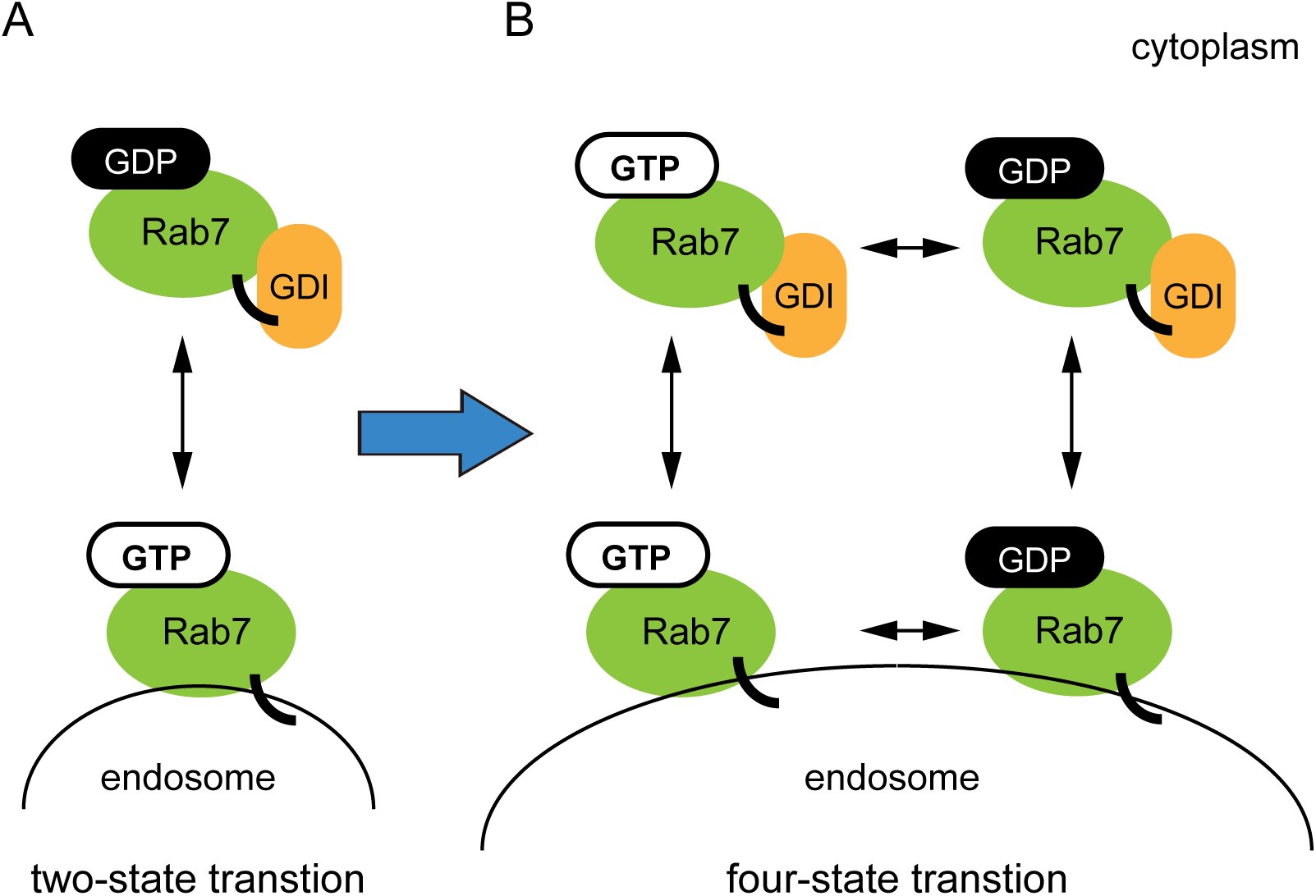
Four-state transition model for Rab7. (A) In a simple two-state model, the prenylated moiety of Rab7-GDP is masked by RabGDI, thereby Rab7-GDP resides in the cytoplasm. On the contrary, the prenylated moiety of Rab7-GTP is unmasked, and thus Rab7-GTP is localized to the endosomal membranes. (B) In a four-state transition model proposed here, both GDP-bound and GTP-bound Rab7 can bind to RabGDI and thus reside in the cytoplasm. Several experimental evidences have shown the existence of Rab7-GDP on endosomal membranes. At present, transitions between respective states are speculative and should be tested experimentally in the near future.

The four-state transition model, particularly the presence of Rab7-GDP on endosomal membranes, has enabled us to account for the discrepancies in experimental data and previously proposed useful but too simplistic “Rab conversion” model (1,14,15). Through a process called Rab conversion, Rab5 on EEs is replaced by Rab7 on LEs/MVBs during early to late endosome maturation (5,6). The complex maturation program of endosomes (from EEs to LEs) exists to close down recycling and other functions of EEs and allow the progression of LEs within the degradative pathway (1). This Rab conversion has been thought to give unidirectionality to the endocytic process. Recent experiments have shown that Mon1-Ccz1 complex and the CORVET-to-HOPS switch play a role in this process (5,16–18). However, a recent immune-electron microscopy study linking the molecular make-up of endosomes with their ultrastructural characteristics (19) indicates that functionally different intermediates of EEs and LEs can be distinguished, and that the transition between EEs and LEs is rather gradual (20). Furthermore, in phagolysosome formation during apoptosis, the overlap between Rab5 and Rab7 was around 60 min (21). We have previously shown that Rab5 and Rab7 reside together on macropinosomes for 5–15 min in EGF-stimulated COS-7 cells (Kawasaki *et al.,* unpublished). Thus, there are many evidences for the formation of a hybrid Rab5/Rab7 compartment (19). Furthermore, some researchers have reported that Rab7 regulates the trafficking between LEs and lysosomes (8, 9, 21), and, at least in some cases, such as EGFR-trafficking in HeLa cells, Rab7 was shown to be dispensable for the delivery of cargos to the LEs (9). It is very difficult to explain these results directly by simple Rab conversion from EEs to LEs. The present study has demonstrated that some fraction of Rab7-GDP exists on endosomal membranes and in EGF-induced macropinocytosis, Rab7 was initially recruited to macropinosomes in a GDP-bound form and subsequently activated during endosome maturation as described above (11).

We believe that these results provide a framework, which naturally extends beyond the two-state Rab conversion model in the case of Rab7; this framework can permit the temporal formation of the hybrid Rab5/Rab7 in endosomes and explains the discrepancies between the localization of dispensable Rab7 on LEs and the hypothetical absence of inactive Rab7 in endosome membranes.

How are the results in this study applicable to other situations? We suppose that the behavior of some Rab GTPases can be better understood based on the four-state transition model. In fact, we have obtained evidences for the existence of Rab35-GDP on membranes and Rab35-GTP in the cytosol (Adachi-Ishido *et al.,* unpublished). Theoretically, it seems to be rather natural postulating the four-state model for Rab GTPases. Efforts to elucidate which state is dominant and which state is relatively minor and transient could prove to be fruitful. What about the time-course of Rab7 activation in other types of endocytosis? In clathrin-dependent endocytosis and phagocytosis, Rab7 has been shown to be essential for the fusion of LEs with lysosomes. Thus, it is plausible that a further activation of Rab7 during fusion between LEs and lysosomes is observed in a range of endocytic processes to accelerate fusion. These issues could be member, cargo, or cell-type specific. Thus, it would be rewarding to create a biosensor for each member of Rab GTPases and examine the changes in its activity in individual contexts. This provides fundamental information to better understand the basic mechanisms regulating Rab machineries implicated in a broad range of physiological and pathological processes.

## Methods

### Plasmids

The cDNAs for wild-type Rab7, Rab7 S22N, and Rab7 Q67L were subcloned into pCAGGS-EGFP (11). The cDNA for wild-type Rab7 was cleaved from EGFP-Rab7 (22) and subcloned into the pBacPac-His3-TEV2bp vector. pGEX-6P-3-RabGDI has been described previously (23).

### Cells, reagents and antibodies

COS-7 cells were maintained in DMEM containing 10% fetal bovine serum. EGF was purchased from Calbiochem. GDP and GTPγS were purchased from Sigma-Aldrich. The following primary antibodies were used: rabbit polyclonal antibodies to Rab7 (Sigma-Aldrich); mouse monoclonal antibody to GST (26H1, Cell Signaling), and anti-GFP polyclonal antibodies, which was a gift from N. Mochizuki.

### Confocal microscopy

Cells were imaged with an IX81 inverted microscope (Olympus) equipped with an FV-1000 confocal imaging system and an automatic programmable XY stage (MD-XY30100T-Meta, SIGMA KOKI) through a plan-apochromat 60× oil objective lens (NA 1.4, Olympus).

### *In vitro* GST pull-down assay

6×His-Rab7 was expressed in Sf9 cells and supplemented with 5 mM mevalonate, a soluble precursor of isopentenylpyrophosphate, which is required for prenylation, in the last 24 h of a 72 h transfection protocol. Cells were then washed with PBS, sonicated four times in buffer A (10 mM Tris-HCl pH 8.0, 1 mM DTT, 10 mM MgCl_2_, and protease inhibitors) and centrifuged at 100,000×g for 1 h. The pellet was resuspended in buffer B (20 mM Tris-HCl pH 8.0, 1 mM DTT, 5 mM MgCl_2_, and 1% CHAPS) and sonicated six times. The suspension was made in 300 mM NaCl and centrifuged as mentioned above. Purification was performed on NTA-resin (Qiagen) and the recombinant protein was eluted in buffer C (20 mM Tris-HCl pH 8.0, 1 mM DTT, 5 mM MgCl_2_, 0.6% CHAPS, and 500 mM imidazole). His-tag was removed by using TEV protease cleavage. GST-RabGDI was prepared as previously described (23). *In vitro* GST pull-down assay were performed as previously described (24) with some modifications. To prepare the GDP and GTPγS bound forms of 6×His-Rab7, buffer 1 (50 mM Tris-HCl, PH 8.0, 2.5 mM DTT, 25 mM EDTA, and 12.5 mM MgCl_2_) containing 20 µM GDP or GTPγS and buffer 2 (20 mM Tris-HCl, pH 8.0, 1 mM DTT, 1 mM EDTA, 5 mM MgCl_2_, and 0.6% CHAPS) containing 6×His-Rab7 were mixed and incubated for 15 min at 30°C. Subsequently, MgCl_2_ at a final concentration of 10 mM was added to this mixture to stabilize the nucleotides. GDP- or GTPγS- loaded 6×His-Rab7 was incubated with GST-RabGDIα and glutathione Sepharose 4B (GE Healthcare Bioscience) for 1.5 h at 4°C and washed three times with buffer 3 (25 mM Tris-HCl, pH 7.5, 150 mM NaCl, 10 mM MgCl_2_, 1% Triton X-100, and 1 mM phenylmethylsulfonyl fluoride). Proteins bound to GST-RabGDI were eluted with SDS-sample buffer. The samples were subjected to SDS-PAGE followed by immunoblotting.

## Acknowledgements

The authors thank Professor Tamotsu Yoshimori and Dr. Keisuke Tabata (Osaka University) for their advice on *in vitro* GST pull-down assay used here. They are grateful for Ms. Kimiko Nakamura for skilled technical assistance and members of the Nakamura Laboratory for their input. This work was supported by grants from JSPS KAKENHI (15K06782) and the Yamada Science Foundation to T.N.

## Author contributions

TN, MF, and GS conceived the study; TN AK, SK, and SM designed and performed experiments; TN, MF, and GS analyzed and interpreted the data; TN wrote the manuscript; all authors discussed the results and commented on the manuscript.

